# Role of intercellular coupling and delay on the synchronization of genetic oscillators

**DOI:** 10.1101/2020.09.29.318717

**Authors:** Supravat Dey, Lee Tracey, Abhyudai Singh

## Abstract

Living cells encode diverse biological clocks for circadian timekeeping and formation of rhythmic structures during embryonic development. A key open question is how these clocks synchronize across cells through intercellular coupling mechanisms. To address this question, we leverage the classical motif for genetic clocks the Goodwin oscillator where a gene product inhibits its own synthesis via time-delayed negative feedback. More specifically, we consider an interconnected system of two identical Goodwin oscillators (each operating in a single cell), where state information is conveyed between cells via a signaling pathway whose dynamics is modeled as a first-order system. In essence, the interaction between oscillators is characterized by an intercellular coupling strength and an intercellular time delay that represents the signaling response time. Systematic stability analysis characterizes the parameter regimes that lead to oscillatory dynamics, with high coupling strength found to destroy sustained oscillations. Within the oscillatory parameter regime we find both in-phase and anti-phase oscillations with the former more likely to occur for small intercellular time delays. Finally, we consider the stochastic formulation of the model with low-copy number fluctuations in biomolecular components. Interestingly, stochasticity leads to qualitatively different behaviors where in-phase oscillations are susceptible to the inherent fluctuations but not the anti-phase oscillations. In the context of the segmentation clock, such synchronized in-phase oscillations between cells are critical for the proper generation of repetitive segments during embryo development that eventually leads to the formation of the vertebral column.

## I. INTRODUCTION

Sustained oscillations are ubiquitous in biological systems from oscillating gene products inside cells to spontaneous beating of cilia and flagella on the cell surface microorganisms and airway epithelium [1]–[3]. Diverse cell types are known to encode genetic oscillators or clocks that function in circadian timekeeping [4], [5], cell cycle regulation [6], [7] and rhythmic structure formations across organisms during development [8]–[10]. An intriguing central question that underlies these phenomena is how cells synchronize oscillation through intercellular coupling processes to create emergent behaviors.

Over the years, several biomolecular motifs have been uncovered for exhibiting sustained oscillations in gene product levels [4], [9]–[13]. Two common regulatory mechanisms used by cells as functional clocks are time-delayed negative feedback, and combined positive-negative feedback circuits [7]. In this paper, we investigate the effect of intercellular coupling on the collective behavior of genetic oscillators where the communication between adjacent cells happens via signaling molecules/pathways. This interaction couples the oscillators in the neighboring cells which leads to synchronization between oscillators. The synchronization of coupled genetic oscillators is a subject of great interest both experimentally [9], [14]–[17] and theoretically [18]–[20]. For examples, in the case of the segmentation clock, coupling of contacting cells through the Notch-Delta signaling pathway leads to perfect formation of vertebral column in vertebrates [10].

Here we study a system of two coupled Goodwin oscillators as illustrated in Fig. 1. The Goodwin oscillator circuit consisting of three species is one of the simplest model based on delayed auto-inhibition that creates sustained oscillations. We phenomenologically model the intercellular coupling via a first-order system that mimics the signaling pathway biochemically connecting the two oscillations. Two essential aspects of the intercellular coupling are preserved in our simplified model– coupling delay and coupling strength. The signaling molecule in one cell represses the gene expression of the other, and the repression strength represents the coupling strength. The response timescale for the first-order dynamics indicates the coupling delay. Using linear stability analysis of the coupled systems, we systematically show that strong coupling can destroy the sustained oscillations, and obtain phase diagram for oscillating and non-oscillating regimes as a function of coupling delay and coupling strength. Then, we characterize the nature of collective oscillations within the oscillatory regime. We find that depending on the value coupling strength and delay the system can show in-phase, anti-phase synchronizations. Finally, we study the synchronization using the stochastic formalism, as biochemical reactions that occur in low molecular copy number subject to inherent randomness. Interestingly, the introduction of stochasticity reduces the region of parameter space for in-phase synchronization.

**Fig. 1.**
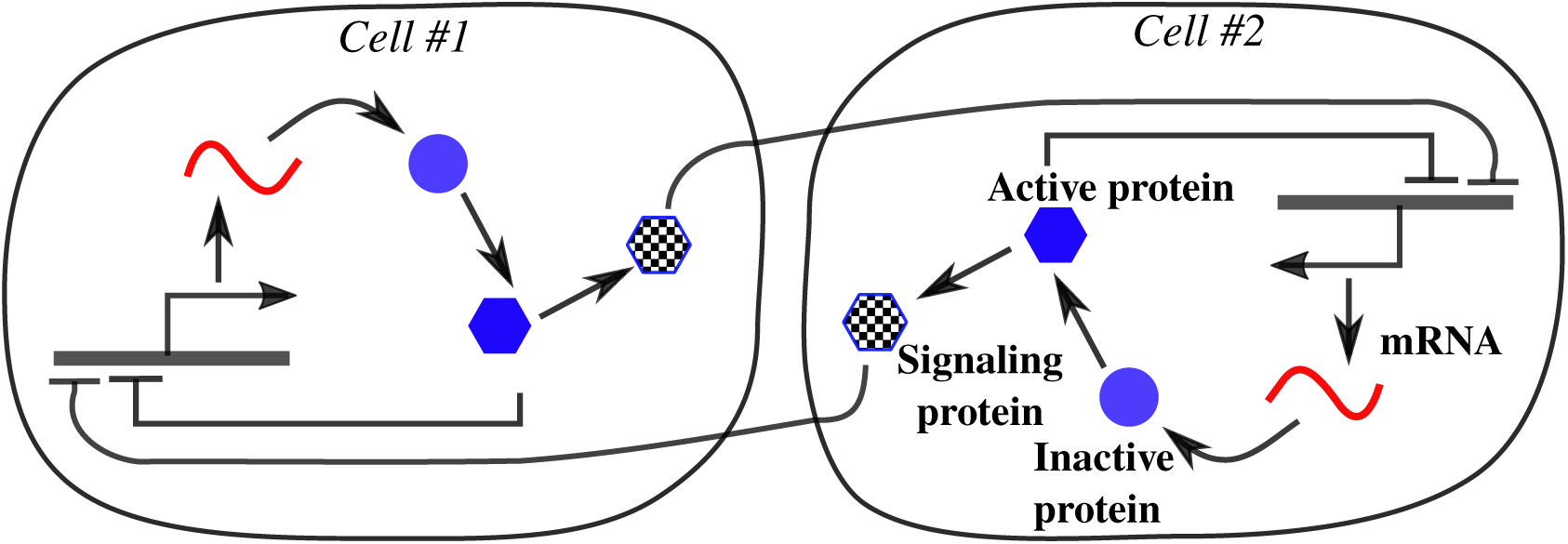
Two Goodwin oscillators coupled via a signaling molecule. Each cell encodes a genetic clock based on the Goodwin oscillator, where a mRNA is translated into the inactive protein. The inactive protein converts into an active protein that represses mRNA synthesis to create a time-delayed negative feedback oscillator. The active protein also activates the production of a signaling molecule that represses mRNA transcription in the neighbouring cell.This simplified coupling is inspired by the segmentation clock where clocks within single cells are coupled by the Notch-Delta signaling pathway to create synchronized traveling waves that generate the vertebral column during development [8]–[10].

## II. Model Formulation

For this study, we consider the well known Goodwin oscillator based on the regulatory mechanism of time delayed auto-inhibition of gene activity [21]. We start by first reviewing the Goodwin oscillator that operates inside an individual cell.

### A. Dynamics of the Goodwin oscillator without coupling

The basic dynamics of a Goodwin oscillator can be represented by the synthesis of a mRNA, that is subsequently translated into an inactive protein. The inactive protein converts into an active form that represses mRNA synthesis to create a delayed negative feedback (Fig. 1). The dynamics of the intracellular concentration of the mRNA (*m*), inactive protein (*p*) and active protein (*x*) evolve as per the nonlinear differential equation

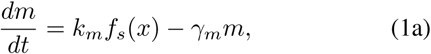

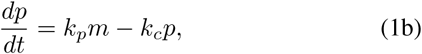

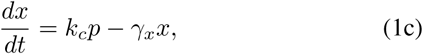

where, *k*_*m*_ is the maximal transcription rate, *k*_*p*_ is the translation rate, *k*_*c*_ is the conversion rate from the inactive protein to active protein, *γ*_*m*_ is the degradation rate of mRNA, and *γ*_*x*_ is the degradation rate for the active protein. The repression function for the single oscillator *f*_*s*_(*x*) is given by

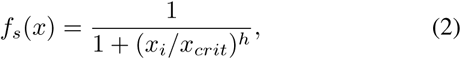

where *h* is the Hill coefficient and *x*_*crit*_ is the concentration where the repression is the half of the maximum value.

Here, we note that to observe sustained oscillations in the context of auto-inhibition and linear degradation, at least an intermediate dynamics as in our case via inactive protein or explicit incorporation of time-delay via delayed differential equation is essential, i.e. a two variables system does not show any sustained oscillations [22]. Below, we discuss the criterion for sustained oscillations for the Goodwin oscillator using linear stability analysis.

#### Linear stability for the single Goodwin oscillator

We linearize the dynamics of single Goodwin oscillator around the fixed point of the system. The fixed point of the system 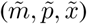 can be obtained by solving the Eq. 1 by setting time derivatives to zero and is given by

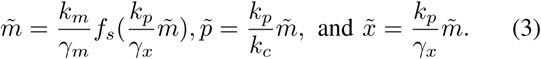

The monotonically decreasing function *f*_*s*_ ensures that the above transcendental equation in 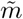 has an unique solution. The matrix for the linearized dynamics of Eq. 1 around the above fixed point is given by the Jacobian matrix

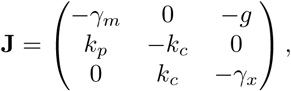

Where

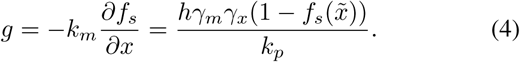

The characteristic polynomial of the Jacobian can be written in the form *Q*(*x*) = *x*^3^ +*ax*^2^ +*bx*+*c*. The polynomial will be stable if (i) *a, b, c >* 0, and (ii) *ab > c* [23]. It can be shown that for the Goodwin dynamics, the first condition always holds true and the violation of the second condition leads to an unstable equilibrium that in this case, manifests as a stable limit cycle. The latter violation implies the following relation

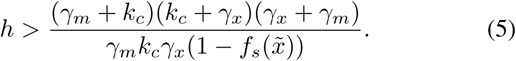

The functional form of the right hand side of the above equation suggests that its minimum value is 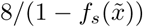, when *k*_*c*_ = *γ*_*m*_ = *γ*_*x*_. Thus, no oscillatory solution exists for the Hill coefficient value less than 8 [22], [24].

### B. Two Goodwin oscillators with intercellular coupling

To incorporate intercellular coupling between two Goodwin oscillators, we introduce two signaling molecules (one for each cell). The signaling molecules is activated by the oscillator and represses gene transcription in the neighboring cell (see Fig. 1). Usually, the signaling molecules takes a finite amount of time to influence (by activating or repressing) the gene expression in the neighbouring cells because it involves several biochemical reactions and the exportation of biomolecules. This involved process in the intercellular coupling is often modeled via a reaction cascade [19] or using an explicit time delay in the dynamical equations [9], [20].

Here, we phenomenologically model the associated delay in the coupling via the first order reaction for the signaling molecule dynamics with an associated timescale *τ* that represents time delay. This signaling molecule then represses the transcription of the neighbouring cell. The coupling is bidirectional and symmetrically acts on both the cells. The dynamics coupled systems is given by

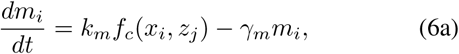

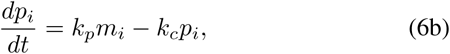

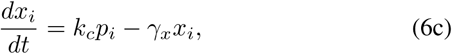

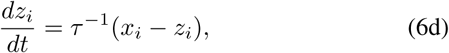

*i* = 1, 2 and *j* is the neighbour of *i*, i.e., *j* = 2(1) for *i* = 1(2). The last equation represents the dynamics of the signaling molecule. The inverse of *τ* is the rate for both the production and degradation of the signaling molecule. The repression functions for the coupled oscillators depends on the level of signaling molecule in the neighbouring cell and we choose it as

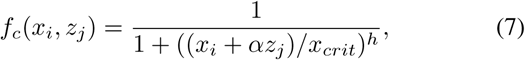

Here *x*_*crit*_ is the concentration of the active protein for which the repression becomes the half of the maximum value in the absence of coupling. The parameter *α* is coupling strength. For no coupling case *α* = 0 and *f*_*c*_ reduces to repression function for the single oscillator *f*_*s*_. On the other hand, *α* = 1 is the maximal coupling strength, assuming the repression due to the intercellular coupling cannot dominate over the intracellular repression due to the active protein.

The above way of incorporating the coupling is motivated by the vertebrate segmentation clock [9], [15], where the active protein represses its own production and also the production of a signaling molecule *deltaC*, and *deltaC* in turn activates the transcription of the gene in in the adjacent cells. We note that *deltaC* should not be confused with the signaling molecule in our systems. In our case, the oscillating gene actives the signaling molecule production not represses as in *deltaC*. However, the repression function together with the signaling molecule dynamics captures the Notch-Delta coupling effectively.

## III. Linear Stability Analysis for Coupled oscillators

Before discussing the effect of coupling on the collective oscillations of the coupled system, we ask the following question: Does the coupling induce or destroy sustained oscillations that were present without coupling? To answer that we do a linear stability analysis of the coupled system. For this, we first determine the fixed points of the dynamical system and then examine the dynamics of the linearized system near the fixed point. The above dynamical system has a unique fixed point given by,

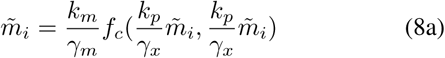

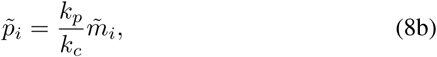

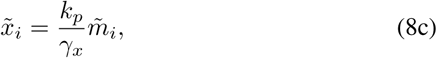

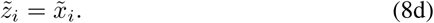

Again as in the case of single Goodwin oscillator, the Eq. 8(a) is a transcendental equation, and the monotonically decreasing function *f*_*c*_ ensures equation has one real solution. It is important to note that the fixed point value depends on the coupling strength *α* but is independent of the coupling delay *τ* in our description.

We now linearize the dynamical equations of the coupled system around the fixed point and the Jacobian matrix is given by,

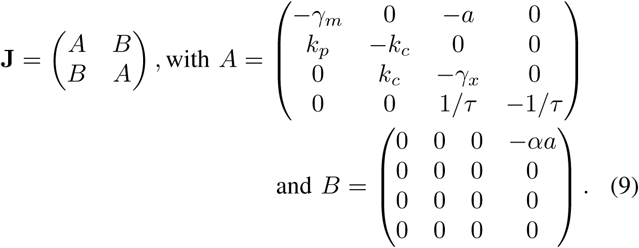

Here, the factor *a* in matrix *A* and *B* is given by,

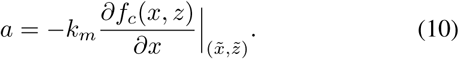

For the coupled system, the order of the characteristic polynomial is 8. Therefore, the obtaining analytical expression of stability criteria for the coupled system is involved unlike for the single Goodwin oscillator where the order of the polynomial is 3. Thus, we obtain the stability condition by numerically looking at the Eigenvalues. We determine the Eigenvalues of the eight dimensional Jacobian matrix for a given set of parameters using Mathematica software (Wolfarm Research, Version 11.3). If the real part of any single Eigenvalue is negative then system is stable and no sustained oscillations can be observed. On the other hand, if the real of any Eigenvalue is positive the dynamics will be unstable where sustained oscillations are observed.

Interestingly, we observe that a high coupling strength can destroy sustained oscillations that are present without coupling. The threshold value of coupling strength above which there are stable oscillations depends on the coupling delay. In Fig. 2(A), we plot phase diagram of the oscillating region as a function of coupling strength *α* and coupling time delay *τ* for a given set of parameter. For small *τ* value, sustained oscillations are observed for any *α*. As *τ* value crosses above a threshold, there are no sustained oscillations above a critical *α* value. Although we have presented the density plot for a given parameter, we have checked that this observation is also holds true for other parameter sets. We numerically solve the ordinary differential equations Eq. (6), and plot the trajectories of mRNA level as a function of time for a given cell in Fig. 2(B). It is clear that sustained oscillations are lost if we change the value of *α* from the oscillatory regime to non-oscillatory regime.

**Fig. 2.**
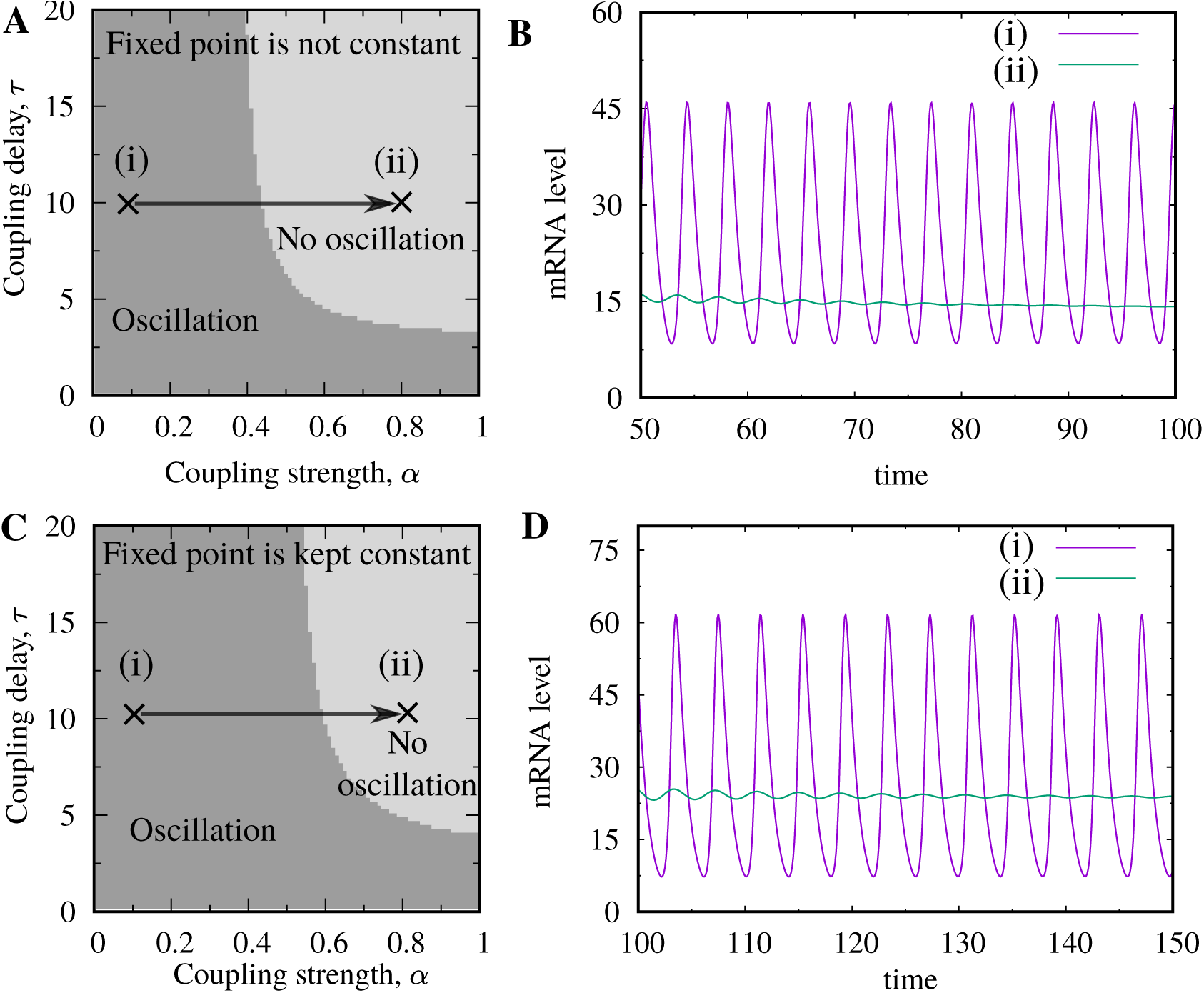
Effect of coupling strength *α* and coupling delay *τ* on oscillating solutions. (Top panel) The fixed point varies with *α*. (A) Above a threshold value of *α* and *τ* a non-oscillatory regime emerges. (B) The trajectory of mRNA for a given cell for representative points in oscillatory (point (i) in subplot (A)) and non-oscillatory (point (ii) in (A)) regime. (Bottom panel) The fixed is kept constant by varying transcription rate *k*_*m*_ with *α* according to Eq. (11). (C) Similar to the top panel a non-oscillatory region is observed for large value of *α* and *τ*. However, the oscillatory region is larger in this case compared to (A). (D) The trajectory of mRNA for a given cell for representative points in oscillatory (point (i) in subplot (C)) and non-oscillatory (point (ii) in (C)) regime. Parameters: *k*_*m*_(0) = 200, *γ*_*m*_ = 1, *k*_*p*_ = 1, *k*_*c*_ = 1, *γ*_*x*_ = 1, *h* = 12, and *x*_*crit*_ = 20.

As we have discussed previously, the change in *α* also affects the fixed point value of the coupled system. Does the loss of sustained oscillations associated with the alteration of the fixed point? To understand this, consider the following case. We vary both *α* and the synthesis rate *k*_*m*_ simultaneously to keep the fixed point value constant to that of without coupling 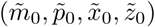.For this, we increase *k*_*m*_ with the *α* in the following way:

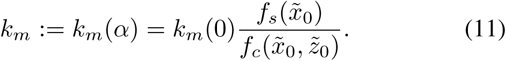

The oscillatory regime for the constant fixed point case is shown in Fig. 2(C). We find the same qualitative behavior with the case when the fixed point is not kept constant (Fig. 2(A)). However, the oscillatory regime becomes larger if we keep the fixed point the same. The plot of typical trajectories mRNA level as a function of time for a given cell for oscillatory and non-oscillatory regime shown in Fig. 2(D). For a given *τ* and *α*, we see that the amplitude of the sustained oscillations in Fig. 2(D) is larger compared to the Fig. 2(B).

## IV. Synchronization of coupled oscillators

The above linear stability analysis cannot reveal whether the same species in both cells oscillate in synchrony or out of synchrony. Below we study the synchronization within the oscillatory region using both the deterministic and stochastic formalisms.

### A. Deterministic Analysis

As the deterministic dynamics is nonlinear (Eq. 6), the collective behavior may depend on the initial conditions of the coupled system. In fact, we have checked that the dynamical response of our system is initial condition dependent. Therefore, to quantify the synchronization, we compute the Pearson correlation coefficient from the steady-state trajectories for large number of random initial conditions. The Pearson correlation between mRNA levels in two cells *C*_12_ is given by,

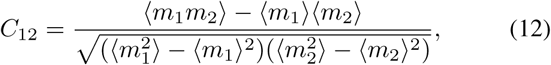

where the angular bracket ⟨·⟩ denotes the average over time and initial conditions. The correlation *C*_12_ takes value 1 if mRNA in both cells oscillate in perfect in-phase, −1 if both cells oscillate in perfect anti-phase, 0 if both cells oscillate randomly.

In Fig. 3(A), we plot *C*_12_ as a function *α* and *τ* for the deterministic case. Depending on the value of *α* and *τ* all the possible scenarios of in-phase, anti-phase, and random-phase oscillations are observed. A clear in-phase oscillation is observed for small values *τ* and large *τ* if *α* is sufficiently large. For the intermediate value of *α*, an anti-phase oscillation is observed when *τ* is higher than a threshold. In between the in-phase and anti-phase regions, the oscillations in both cells are random more or less. In Fig. 3(B) and Fig. 3(C), we plot the trajectories the mRNA level for both cells for the representative points (i) (large *α* and small *τ*) and (ii) (intermediate *α* and small *τ*) marked in the density plot (Fig. 3(A)) for a random initial condition. In-phase is seen when for a small *τ*. Whereas, the anti-phase synchronization is observed for large *τ* for an intermediate value of *α*.

**Fig. 3.**
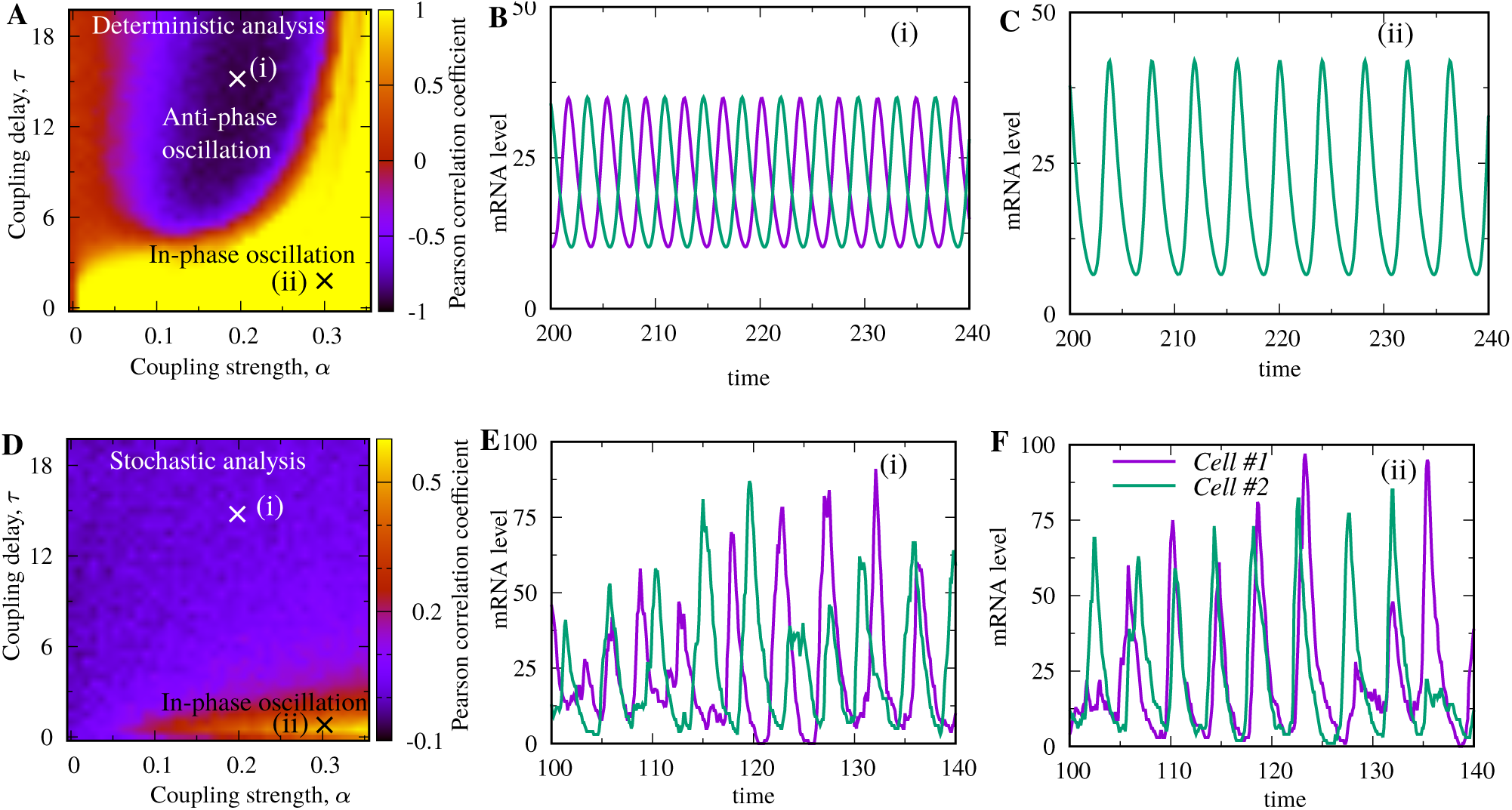
Effect of coupling strength and time delay on the synchronization between two Goodwin oscillators. (Top panel) Deterministic case: (A) Density plot for the Pearson correlation coefficient between mRNA levels of both cells, *C*_12_, as a function of *α* and *τ* showing in-phase (*C*_12_ *>* 0), anti-phase (*C*_12_ *<* 0) and random-phase (*C*_12_ ≈0) regions. *C*_12_ is are averaged over 400 initial conditions. The points (i) and (ii) (marked by the cross symbol) are two representative points showing in- and anti-phase synchronizations. The trajectories of mRNA levels for both cells for a given initial condition are shown in (B) and (C) for the point (i) and (ii), respectively. (Bottom panel) Stochastic case: (D) Density plot *C*_12_ as a function of *α* and *τ*. The region of in-phase synchronization becomes quite small compared to the deterministic case. There is no prominent region of anti-phase synchronization. The trajectories for two representative points (i) and (ii) for this stochastic case are shown in (E) and (F). The stochastic trajectory for both the cells oscillate in synchrony for (ii). While the anti-phase synchronization between the stochastic trajectory for both cells is not clear for (i) as observed in the deterministic case. Parameters: *k*_*m*_(0) = 200, *γ*_*m*_ = 1, *k*_*p*_ = 1, *k*_*c*_ = 1, *γ*_*x*_ = 1, *h* = 12, and *x*_*crit*_ = 20. The fixed point is not kept constant by varying *k*_*m*_.

### B. Stochastic Analysis

So far, we have been discussing the deterministic dynamics of the coupled Goodwin oscillators. However, the gene expression is stochastic due to underlying biochemical reactions that are inherently stochastic [25]–[31]. Besides, there are extrinsic perturbations (due to factors such as cell-to-cell differences in expression machinery) that can make an oscillatory expression noisy [29], [32]–[38]. As a result simple non-oscillatory gene expressions [25]–[30] as well oscillatory gene expressions [10], [39], [40] are subject to large fluctuations. The stochastic oscillatory gene expression can have several implications on biological functions and the control of stochasticity is desire in many cases [17], [24], [41], [42]. Therefore, it is importantto study the synchro-nization study using stochastic formalism.

In stochastic the setting, biochemical reactions occur randomly. After each reaction occurs, the number of involved species change discretely. We denote molecular count of the mRNA, inactive protein, active protein, and signaling molecule in cell *i* at time *t* by *m*_*i*_(*t*), *p*_*i*_(*t*), *x*_*i*_(*t*), and *z*_*i*_(*t*), respectively. The transition probabilities of all the reactions within an infinitesimal time interval *t* and *t* + *dt* in cell *i* ∈ {1, 2} are given by

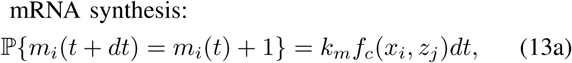

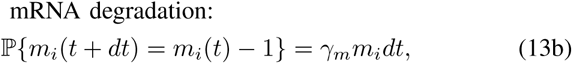

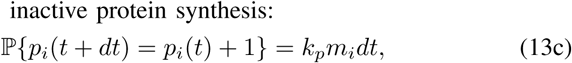

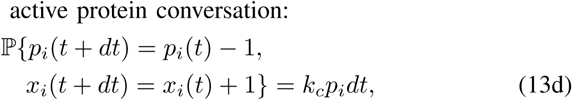

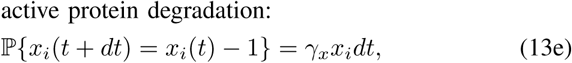

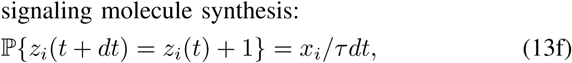

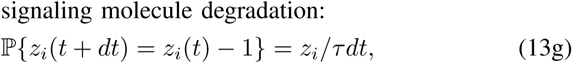

where *j* is the neighbor of cell *i*.

We use the Gillespie algorithm to solve the stochastic dynamics numerically and generate stochastic trajectories [43]. The typical stochastic trajectories mRNA level for both both cells are shown in Fig. 3(E) and (F). Using a large number of stochastic trajectories, we compute the Pearson correlation coefficient *C*_12_ as we did in the deterministic analysis, here, the angular bracket ⟨·⟩ in Eq. 12 represents the average of time and ensembles. We plot *C*_12_ in Fig. 3(D) as a function of *α* and *τ*. The plot is not the same as the deterministic case. The parameter regime where oscillations show in-phase becomes very small compared to the deterministic case the anti-phase synchronization is not prominent. This reduction of parameter space for synchronized oscillations suggests that inherent stochasticity can destroy the established correlation between two cells due to the coupling. The loss of correlation also is also clear from the trajectory plots (Fig. 3(E) and (F)).

## V. Conclusion and discussion

In summary, we have investigated the role of intercellular coupling on the collective oscillations between two Goodwin oscillators. Motivated by real Delta-Notch signaling, we have phenomenologically modeled the intercellular coupling between cells via the first-order dynamics of a repressive signaling molecule. While the repression strength corresponds to the coupling strength, the associated timescale of the first-order dynamics represents time delay in coupling.

We have found both the coupling strength and coupling delay play a crucial role in the coupled dynamics. Using linear stability analysis, we have shown that large coupling can destroy sustained oscillations if the coupling delay is larger than a threshold. We have found that, depending on the value of the coupling strength and delay, that the species in both cells can oscillate in-phase synchrony, anti-phase synchrony or randomly. We note that the appearance of alternating multiple in-phase and anti-phase was previously reported where the intercellular coupling was modeled via a reaction cascade [19] or incorporating delay in explicitly in the deferential equation [20]. However, in our system, we do not see multiple appearance of in-phase and anti-phase.

Finally, as the species levels in real oscillators are subject to large fluctuations due to inherent randomness of biochemical reactions, we have studied the correlated dynamics using stochastic formalism. We have found stochasticity can destroy the correlation established by the intercellular coupling. The parameter space for in-phase oscillations become smaller compared to the deterministic case. The impact of stochasticity on anti-phase synchronization is more drastic. We found the anti-phase is not susceptible to inherent stochasticity. We note that in the context of a previous study where the intercellular coupling was modeled via a reaction cascade, both the in-phase and anti-phase found to be stable [19].

## ACKNOWLEDGMENT

This work is supported by the National Science Foundation Grant ECCS-1711548.

